# Cardiac Gq receptors and calcineurin activation are not required for the hypertrophic response to mechanical left ventricular pressure overload

**DOI:** 10.1101/2020.12.08.393595

**Authors:** Ze-Yan Yu, Hutao Gong, Jianxin Wu, Yun Dai, Scott H Kesteven, Diane Fatkin, Boris Martinac, Robert M Graham, Michael P Feneley

**Author notes:** Correspondence to: Michael P Feneley MD, Cardiology Department, St Vincent’s Hospital, 390 Victoria Street, Darlinghurst, NSW 2010, Australia. These authors contributed equally (Ze-Yan Yu, Hutao Gong).

## Abstract

**Rationale:** Gq-coupled receptors are thought to play a critical role in the induction of left ventricular hypertrophy (LVH) secondary to pressure overload, although mechano-sensitive channel activation by a variety of mechanisms has also been proposed, and the relative importance of calcineurin- and calmodulin kinase II (CaMKII)-dependent hypertrophic pathways remains controversial.

**Objective:** To determine the mechanisms regulating the induction of LVH in response to mechanical pressure overload.

**Methods and Results:** Transgenic mice with cardiac-targeted inhibition of Gq-coupled receptors (GqI mice) and their non-transgenic littermates (NTL) were subjected to neurohumoral stimulation (continuous, subcutaneous angiotensin II (AngII) infusion for 14 days) or mechanical pressure overload (transverse aortic arch constriction (TAC) for 21 days) to induce LVH. Candidate signalling pathway activation was examined. As expected, LVH observed in NTL mice with AngII infusion was attenuated in heterozygous (GqI^+/-^) mice and absent in homozygous (GqI^-/-^) mice. In contrast, LVH due to TAC was unaltered by either heterozygous or homozygous Gq inhibition. Gene expression of atrial natriuretic peptide (ANP), B-type natriuretic peptide (BNP) and α-skeletal actin (α-SA) was increased 48 hours after AngII infusion or TAC in NTL mice; in GqI mice, the increases in ANP, BNP and α-SA in response to AngII were completely absent, as expected, but all three increased after TAC. Increased nuclear translocation of nuclear factor of activated T-cells c4 (NFATc4), indicating calcineurin pathway activation, occurred in NTL mice with AngII infusion but not TAC, and was prevented in GqI mice infused with AngII. Nuclear and cytoplasmic CaMKIIδ levels increased in both NTL and GqI mice after TAC but not AngII infusion, with increased cytoplasmic phospho- and total histone deacetylase 4 (HDAC4) and increased nuclear myocyte enhancer factor 2 (MEF2) levels.

**Conclusion:** Cardiac Gq receptors and calcineurin activation are required for neurohumorally mediated LVH but are not required for LVH induced by mechanical pressure overload (TAC); the latter is mediated by activation of the CaMKII-HDAC4-MEF2 pathway.

## Introduction

Pathological left ventricular hypertrophy (LVH) is the single strongest predictor of cardiovascular mortality and heart failure(Levy et al., 1990;Mudd and Kass, 2008), stimulating extensive research efforts to identify the molecular signalling pathways responsible for its induction as potential therapeutic targets(Mudd and Kass, 2008;Tamargo and Lopez-Sendon, 2011). Gq-coupled receptors are thought to play a central role in the induction of pathological LVH in response to agonists, such as angiotensin II (AngII), catecholamines and endothelin-1, or increased mechanical stress associated with hypertension, aortic stenosis and pathological ventricular remodelling after myocardial injury (D’Angelo et al., 1997;Adams et al., 1998;Akhter et al., 1998;Paradis et al., 2000;Frey and Olson, 2003).

At the sub-receptor level, an increase in intracellular calcium and activation of calcium-calmodulin dependent pathways has been thought to be necessary for the induction of pathological LVH(Bers, 2008;Zarain-Herzberg et al., 2011). Activation of the calcium-calmodulin dependent phosphatase, calcineurin, dephosphorylates the transcriptional regulator, nuclear factor of activated T cells (NFAT), resulting in its translocation to the nucleus to initiate hypertrophic gene transcription in pathological but not physiological hypertrophy(Molkentin et al., 1998;Molkentin, 2000;Wilkins et al., 2004;Wilkins and Molkentin, 2004;Kehat and Molkentin, 2010;Molkentin, 2013). Activation of calcium-calmodulin dependent kinase II (CaMKII) phosphorylates histone deacetylase 4 (HDAC4), promoting its nuclear export and relieving its inhibition of the critical nuclear transcriptional regulator, myocyte enhancer factor 2 (MEF2), thus enhancing hypertrophic gene transcription(Passier et al., 2000;Backs et al., 2006;Backs et al., 2009). Both the calcineurin-NFAT and the CaMKII-HDAC-MEF2 pathways have been claimed to be both sufficient and *necessary* for the induction of pathological LVH, although not without considerable controversy(Ding et al., 1999;Zhang et al., 1999;Molkentin, 2000;Zou et al., 2001;Frey and Olson, 2003;Zhang et al., 2003;Zhang et al., 2005;Zhang et al., 2007;Ling et al., 2009;Molkentin, 2013). The question of the relative importance of calcineurin and CaMKII activation in pathological hypertrophy remains under active investigation(Kreusser et al., 2014;Lehmann et al., 2018).

Transient receptor potential canonical (TRPC) channels have been identified as a likely mechanism by which calcium enters through the myocyte cell membrane to trigger calcium-dependent activation of LVH signalling pathways(Watanabe et al., 2009;Wu et al., 2010;Eder and Molkentin, 2011). While activation of Gq-coupled receptors by agonists or mechanical deformation is thought to cause receptor-operated TRPC channel activation by generating diacylglycerol (DAG), and also increased TRPC expression, some TRPC channels can be directly activated by mechanical deformation (Bush et al., 2006;Kuwahara et al., 2006;Onohara et al., 2006;Spassova et al., 2006;Ohba et al., 2007;Seth et al., 2009;Koitabashi et al., 2010;Eder and Molkentin, 2011). TRPCs might also promote calcium entry by functional interaction with other calcium entry mechanisms, including the sodium-calcium exchanger, Orai or L-type calcium channels (Eder and Molkentin, 2011). Alternatively, L-type calcium channels have been strongly implicated as a source of the triggering calcium by a different, protein-binding mechanism that anchors calcineurin to create a microdomain with L-type calcium channels at the sarcolemma(Heineke et al., 2010). Ca(v)3.2 T-type calcium channels (T-channels) have also been claimed to be required for pressure overload-induced cardiac hypertrophy in mice (Chiang et al., 2009). Syndecan-4, a transmembrane proteoglycan that connects extracellular matrix proteins to the cardiomyocyte cytoskeleton, has been reported to be necessary to directly mediate calcineurin activation in response to mechanical deformation and pressure overload (Finsen et al., 2011).

Thus, it is now apparent that there are a variety of mechanisms whereby pathological LVH could be triggered that do not necessarily depend on activation of cardiac Gq-coupled receptors. These mechanisms may be more important when mechanical loading is the predominant stimulus to LVH rather than neurohumoral activation. It is timely, therefore, to revisit the role of Gq-coupled receptors in the induction of LVH in response to left ventricular pressure overload. We show here that in mice with cardiac-targeted transgenic inhibition of Gq(Akhter et al., 1998), LVH in response to continuous AngII infusion was markedly inhibited in heterozygous mice and completely abolished in homozygous mice, as expected. In contrast, heterozygous or homozygous inhibition of Gq had *no effect* on LVH induced by LV pressure overload caused by transverse aortic arch constriction (TAC), the most commonly employed experimental model of pressure overload LVH (Rockman et al., 1994;Frey and Olson, 2003;Molkentin, 2013). Moreover, while AngII infusion caused activation of the calcineurin-NFAT pathway, as expected, it did not activate the CaMKII-HDAC4-MEF2 pathway. Conversely, TAC activated the CaMKII-HDAC4-MEF2 pathway but did not activate the calcineurin-NFAT pathway.

These findings have significant implications for the mechanistic understanding of LVH secondary to pressure overload but may also have important clinical and therapeutic implications for the optimal management of hypertensive heart disease.

## Methods

### Generation of transgenic mice

Transgenic mice with endogenous, cardiac-restricted inhibition of Gq receptors (GqI mice; kindly provided by Dr. W. Koch), described previously in the heterozygous state (GqI^+/-^) (Akhter et al., 1998), were employed. To ensure robust endogenous inhibition of cardiac Gq receptors, homozygous (GqI^-/-^) mice were also bred. Male GqI^-/-^, GqI^+/-^ and non-transgenic littermate (NTL) mice aged 12 weeks were used. All experimental procedures were approved by the Garvan/St Vincent’s Hospital Animal Ethics Committee, in accordance with the guidelines of the Australian Code of Practice for the Care and Use of Animals for Scientific Purposes. All animals were entered into the study in a randomized order.

### Induction of left ventricular hypertrophy

Transgenic and NTL mice were subjected to either continuous subcutaneous infusion of the Gq-coupled receptor agonist AngII or transverse aortic arch constriction (TAC) to induce pressure overload.

For AngII or saline infusion, mice were anaesthetised with 1.5% - 2% isoflurane, and a small incision was made in the skin between the scapulae. An osmotic mini-pump (Alzet) was inserted subcutaneously, and the incision closed. Angiotensin II (1.5 mg/kg/day, Sigma, Australia) was dissolved in 0.9% NaCI and delivered continuously until sacrifice at 48 hours or 14 days. This dose was selected after dose-response testing to find the minimum dose necessary to produce a mild increase in systolic pressure (≈25 mmHg) for 14 days treatment, with the aim of achieving primarily humoral stimulation of LVH. Control mice underwent the same procedure with mini-pumps filled with 0.9% NaCI only.

For TAC, mice were anesthetized with 5% isoflurane and ventilated at 120 breaths/min (Harvard Apparatus Rodent Ventilator). The transverse aortic arch was accessed via an incision in the second intercostal space, and constricted with a ligature tied around a 25-gauge needle between the left and the right arteries, which was then removed (Rockman et al., 1994). TAC does not activate the renin-angiotensin system (Wiesner et al., 1997). Sham mice underwent the same procedure but the ligature was not tied. Simultaneous direct pressure recordings (1.4 F pressure catheter, AD Instruments, P/L) from both the right carotid artery and the aorta distal to the ligature (n=20 mice) indicated a TAC pressure gradient of 60±8 mmHg with this technique. Animals were sacrificed after 48 hours or 21 days.

### Invasive hemodynamic measurements

After 14 days of AngII infusion or 21 days of TAC, mice were anesthetized by inhalation of isoflurane (1.5%) and a 1.4F micro-tip pressure catheter (Millar Instruments Inc, Houston, Texas, USA) was inserted into the LV via the right carotid artery. The heart rate, systolic aortic pressure, LV systolic pressure, +dP/dt, and –dP/dt were recorded (LabChart 6 Reader, AD Instruments, P/L). Animals were sacrificed, and the LV weights (LVW) normalized to body weight (BW) and to tibial length (TL) were measured as indicators of LVH.

### Quantitative Real-Time Polymerase Chain Reaction (RT-PCR)

Gene expression was determined by quantitative RT-PCR. Total RNA was extracted and purified from LV tissue with a QIAGEN RNeasy Fibrous Tissue Mini Kit (QIAGEN, Cat#74704). RNA (500 ng) was reverse transcribed into cDNA using SuperScript III First-Strand Synthesis SuperMix system (Invitrogen, Cat#11752-250). cDNA was subjected to PCR amplification to detect ANP, BNP, α-SA and myocyte-enriched calcineurin-interacting protein 1 (MCIP1 [RCAN1]) gene expression, quantified by the LightCycler 480 Probes master (Roche, Cat#047049001). Probes for quantititative PCR were purchased from ABI: hypoxanthine-guanine phosphoribosyl transferase 1 (HPRT) (Mm00446968.m1), ANP (Mm01255748_-_g1), BNP (Mm01255770_-_g1), α-SA (Acta1, Mm00808218.g1) and MCIP1.4 (Mm1213407__-_m1). Samples were run in triplicate and mRNA levels were normalised to those of HPRT.

### Western blotting

LV tissue was lysed using NE-PER nuclear and cytoplasmic extraction reagents (Pierce Biotechnology, Rockford, Il, USA) and Protease Inhibitor Cocktail Kit and Halt Phosphatase Inhibitor Cocktail (Pierce Biotechnology), homogenised and proteins quantified using the Pierce BCA Protein Assay Kit. Protein (40 μg) was separated by SDS-PAGE, transferred to PVDF membranes (Bio-Rad Laboratories), and blocked with 5% bovine serum albumin (BSA, Sigma).

Primary antibodies included: NFATc4 (1:1500 final dilution; Abcam), phosphorylated (p-) GSK3β (Ser9, 1:1500; Cell Signalling), total GSK3β (1:500; Cell Signalling), GATA4 (1:1000; Santa Cruz Biotechnology), CaMKIIδ (1:1000; Santa Cruz Biotechnology), p-HDAC4 (1:1500; Cell Signalling), total HDAC4 (1:1500; Cell Signalling), MEF2A (1:3000; Cell Signalling). Glyceraldehyde-3-phosphate dehydrogenase (GAPDH, 1:5000; Abcam) and histoneH2B (1:5000; Abcam) were used to standardize for loading. Horseradish peroxidase-conjugated goat anti-mouse (1:5000) or anti-rabbit (1:10000) secondary antibodies (Abcam, MA) were used at room temperature for 1 hour. Immunologic detection was accomplished using Amersham ECL Western blotting detection reagents (GE Healthcare). Protein levels were quantified by densitometry using NIH ImageJ analysis software. Protein levels were normalised to relative changes in histoneH2B for the nuclear fraction (Histone) and GAPDH for the cytoplasmic fraction and expressed as fold changes relative to those of control animals.

### Statistical analysis

All experiments and analyses were blinded. Data are presented as mean ± SEM. Paired 2-tailed Student’s test was applied to determine the statistical significance between 2 groups. One-way ANOVA was used to compare differences among multiple groups, followed by Tukey’s post-hoc test for significance. P < 0.05 was considered significant.

## Results

### Inhibition of cardiac Gq-coupled receptors causes no abnormal cardiac phenotype under baseline conditions

To ensure robust endogenous inhibition of cardiac Gq receptors, we bred not only heterozygous (GqI^+/-^) but also homozygous (GqI^-/-^) mice. GqI^+/-^ and GqI^-/-^ mice at 12 weeks of age were healthy, with no apparent cardiac morphological or pathological abnormalities compared with their NTLs, suggesting that complete inhibition of cardiac Gq receptors did not appreciably alter the baseline phenotype of the heart.

### Inhibition of cardiac Gq-coupled receptors inhibits the induction of left ventricular hypertrophy in response to AngII infusion

We next examined the role of cardiac Gq receptors in the induction of LVH in response to either AngII infusion (a predominantly humoral stimulus) or TAC (a mechanical stimulus) in transgenic (GqI^+/-^ and GqI^-/-^) and NTL mice.

Angiotensin II infusion for 14 days induced significant LVH in NTL mice, evidenced by a 31% increase in LVW/BW and a 28% increase in LVW/TL (p<0.001, Figure 1A). As expected, the degree of LVH induced by AngII was significantly reduced in GqI^+/-^ mice (19% and 17% increases in LVW/BW and LVW/Tl, respectively, p<0.001), and LVH was prevented in GqI^-/-^ mice (Figure. 1A).

**Figure 1:**
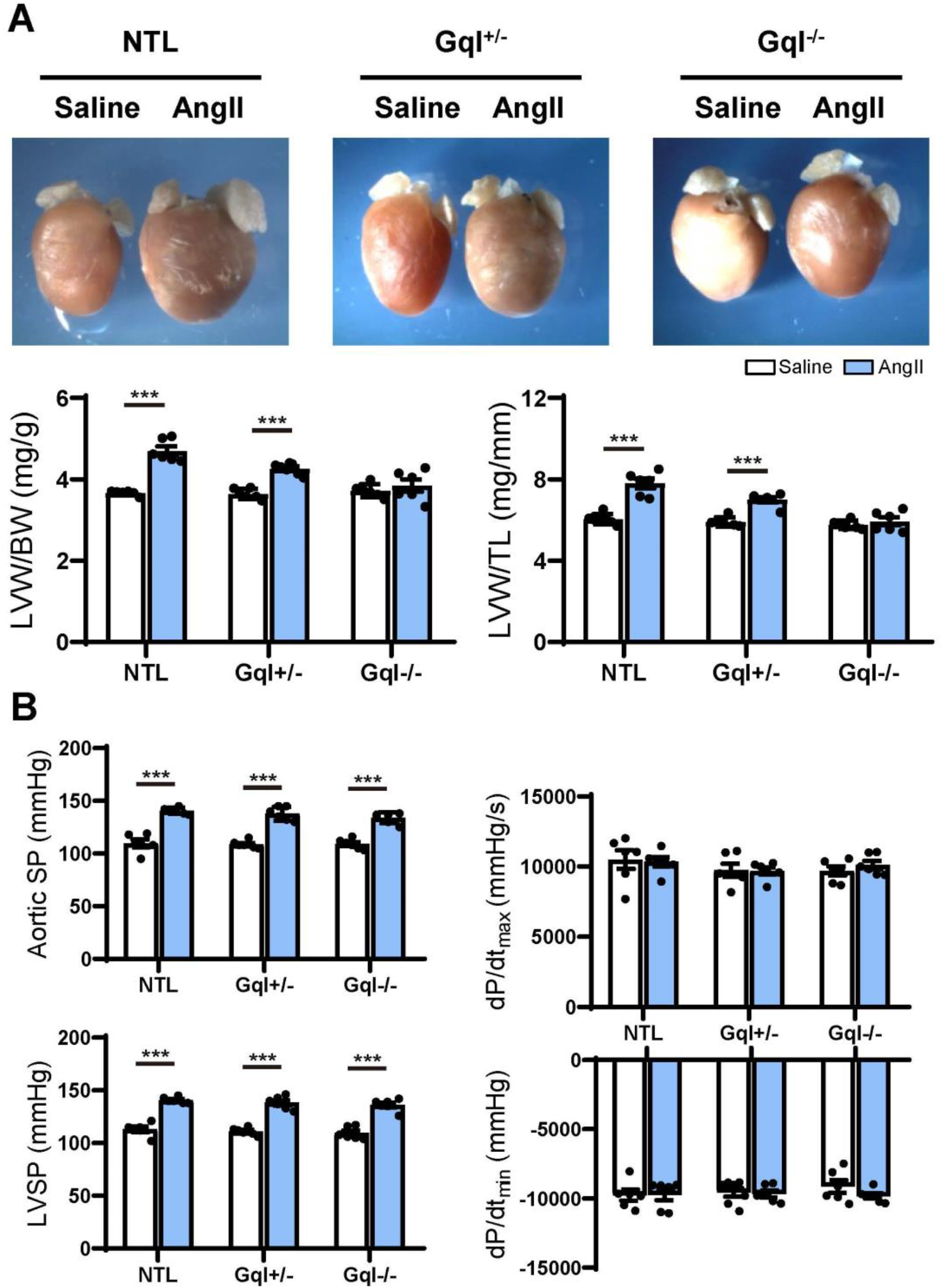
The hypertrophic response to angiotensin II (AngII) infusion after 14 days. (A) Representative hearts and developed LVH indexed by the ratios of LVW/BW and LVW/TL in mice treated with AngII infusion versus saline infusion controls (n=7-9/group); (B) Hemodynamic measurements. All values are mean ± SEM. \*\*\**p*<0.001 vs. saline-treated controls; NS: not significant. LVW/BW: LV weight to body weight ratio; LVW/TL: LV weight to tibia length ratio; SP: systolic pressure; dP/dt: first derivative of pressure with respect to time.

The aortic systolic pressure was 20 mmHg higher, on average, with AngII infusion than with saline infusion (Figure 1B), indicating the small pressor effect of the dose of AngII selected (see Methods), but this effect was the same in NTL, GqI^+/-^ and GqI^-/-^ mice. The complete abolition of the hypertrophic response to AngII in GqI^-/-^ mice despite the same pressor effect seen in NTL mice confirms that humoral activation of the Gq-coupled AngII type 1 (AT1) receptor by AngII was the major mechanism of LVH induction, as distinct from a secondary effect of the small pressor response. There were no significant differences in dP/dt_max_ or dP/dt_min_ between mice treated with AngII or saline in transgenic or NTL mice (Figure 1B).

### Inhibition of cardiac Gq-coupled receptors has no effect on the induction of left ventricular hypertrophy in response to TAC

After 21 days, TAC induced very significant LVH in NTL mice, evidenced by a mean 56% increase in LVW/BW and a mean 48% increase in LVW/TL relative to sham-operated animals (both p<0.001, Figure 2A). Remarkably, the hypertrophic response to TAC observed in NTL mice was the *same* in GqI^+/-^ and GqI^-/-^ mice, evidenced by 58% and 61% mean increases, respectively, in LVW/BW and 52% and 51% mean increases, respectively, in LVW/TL (all p<0.001, Figure 2A).

**Figure 2:**
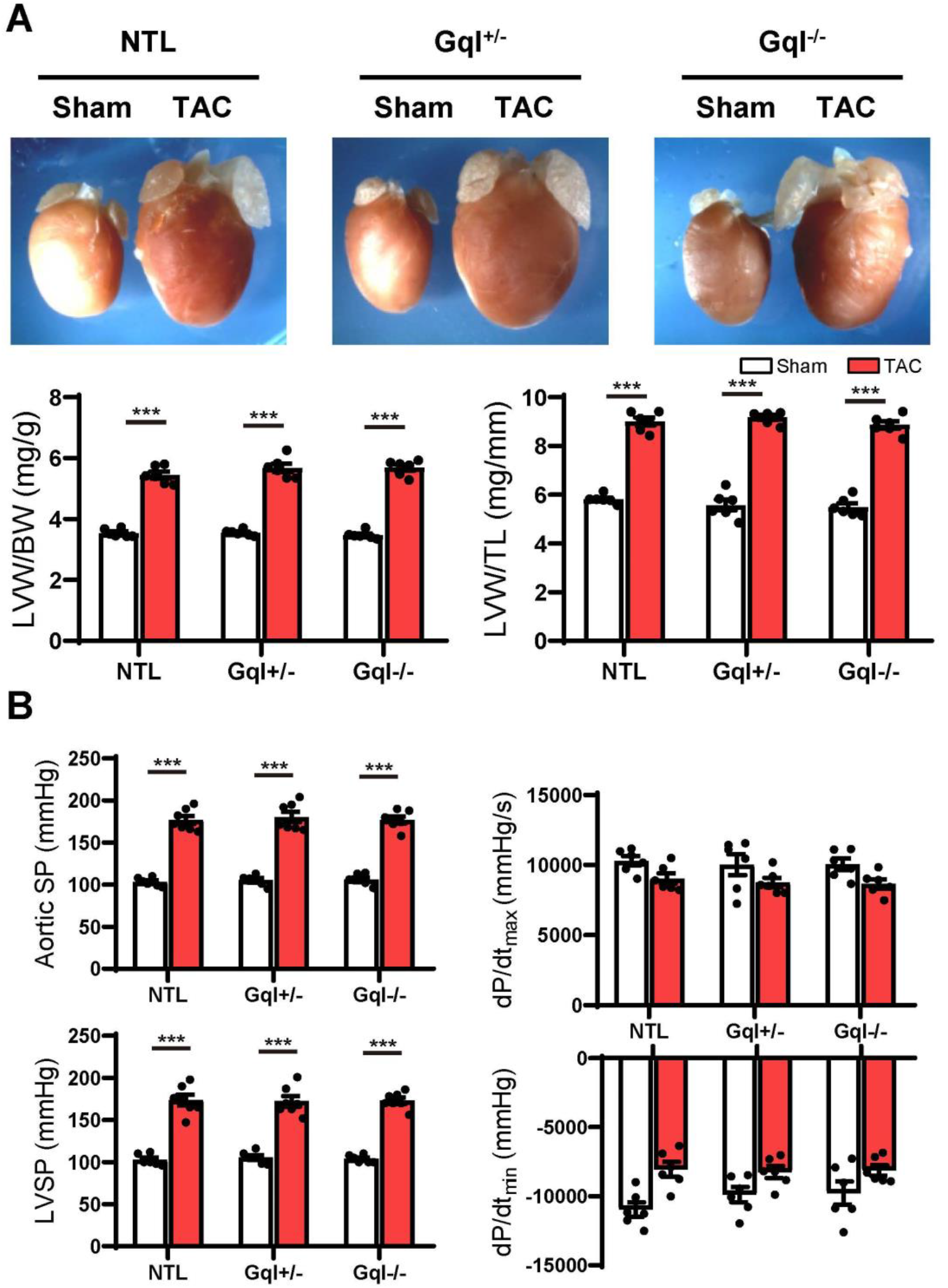
The hypertrophic response to TAC for 21 days. (A) Representative hearts (upper) and developed LVH indexed by the ratios of LVW/BW and LVW/TL (lower) in mice subjected to TAC versus sham-operated controls (n=7-9/group); (B) Hemodynamic measurements. All values are mean ± SEM. \*\**p*<0.01, \*\*\**p*<0.001 vs. sham-operated controls; LVW/BW: LV weight to body weight ratio; LVW/TL: LV weight to tibia length ratio; SP: systolic pressure; dP/dt: first derivative of pressure with respect to time.

Aortic pressure proximal to the constriction and LV systolic pressure increased by 60-70 mmHg with TAC (Figure 2B). Importantly, the increase in the pressure load on the LV with TAC was the same in NTL, GqI^+/-^ and GqI^-/-^ mice. A statistically non-significant and equal reduction in dP/dt_max_ and dP/dt_min_ was observed with TAC in all three groups. There was no difference in lung weight normalised to body weight in TAC mice and sham-operated mice in any of the three groups (data not shown), indicating that this TAC model at 21 days remains a model of compensated LVH rather than heart failure.

### Early markers of induction of left ventricular hypertrophy

Although there was no significant LVH after 48 hours of either AngII infusion or TAC, as indexed by the ratios of LVW/BW or LVW/TL, the early induction of hypertrophic pathways was evidenced by dramatic (3-8 fold) increases in gene expression of ANP, BNP and α-SA in NTL mice with both AngII and TAC (Figure 3). In contrast, in both GqI^+/-^ and GqI^-/-^ mice, there was no increase in gene expression of ANP, BNP or α-SA with AngII infusion, but the increases in gene expression of ANP, BNP and α-SA with TAC in transgenic GqI mice were similar to those observed in NTL mice. These results mirror those described above regarding LVH measured after 14 days or 21 days of AngII infusion or TAC, respectively.

**Figure 3:**
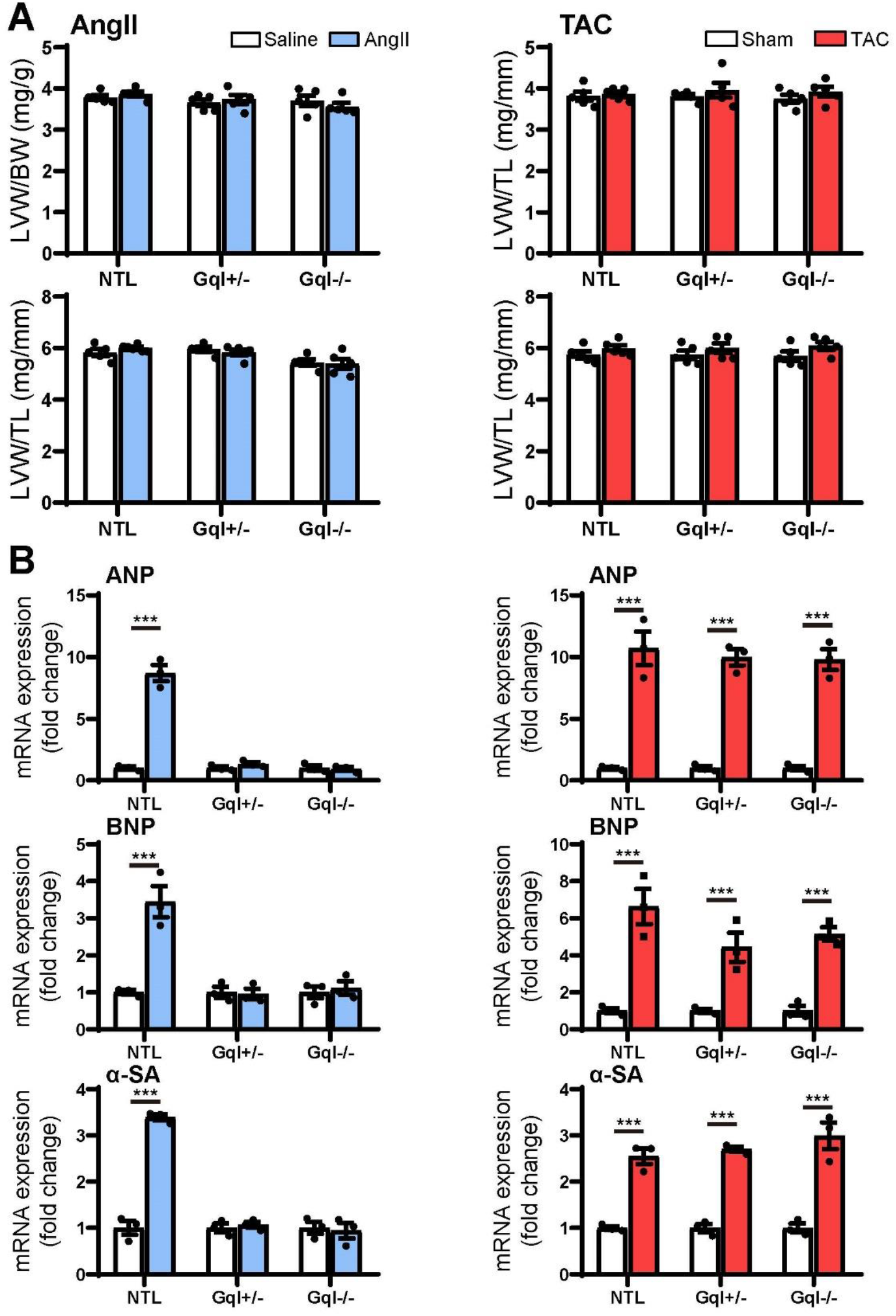
Markers of the early induction of hypertrophy in response to angiotensin II infusion (AngII) (left panels) or TAC (right panels) after 48 hours. (A) Ratios of LVW/BW and LVW/TL, n=6-8/group. (B) Gene expression of ANP, BNP and α-skeletal actin, n=4-5/group. All values are mean ± SEM. \*\*\**p*<0.001 vs. saline treated or sham-operated controls, respectively; NS: not significant. LVW/BW: LV weight to body weight ratio; LVW/TL: LV weight to tibia length ratio; ANP: atrial natriuretic peptide; BNP: B-type natriuretic peptide; α-SA: α-skeletal actin.

### Angiotensin II infusion activates the calcineurin-NFAT hypertrophic signaling pathway, and this activation is inhibited by inhibition of cardiac Gq-coupled receptors

Infusion of AngII for 48 hours in the NTL mice led to increased NFATc4 translocation to the nucleus, evidenced by a 66% mean increase in nuclear NFATc4 (*p*<0.01) without change in cytoplasmic NFATc4, with a consequent 68% mean increase in the nuclear/cytoplasmic NFAT ratio (p<0.01) when compared with saline-treated controls (Figure 4). These data indicate calcineurin activation by AngII infusion. The increase in NFATc4 nuclear translocation in NTL hearts was accompanied by a significant increase in the level of the nuclear transcription factor GATA4 (*p*<0.05). Calcineurin activation by AngII infusion in NTL hearts was also accompanied by an increase in the nuclear fraction of the serine 9 phosphorylated form of glycogen synthase kinase-3 beta (GSK3β, *p*<0.01) but no change in total nuclear GSK3β and no change in the total or phosphorylated form of GSK3β in the cytoplasm. These data are consistent with calcineurin’s inhibition of the nuclear export of NFAT by GSK3β.

**Figure 4:**
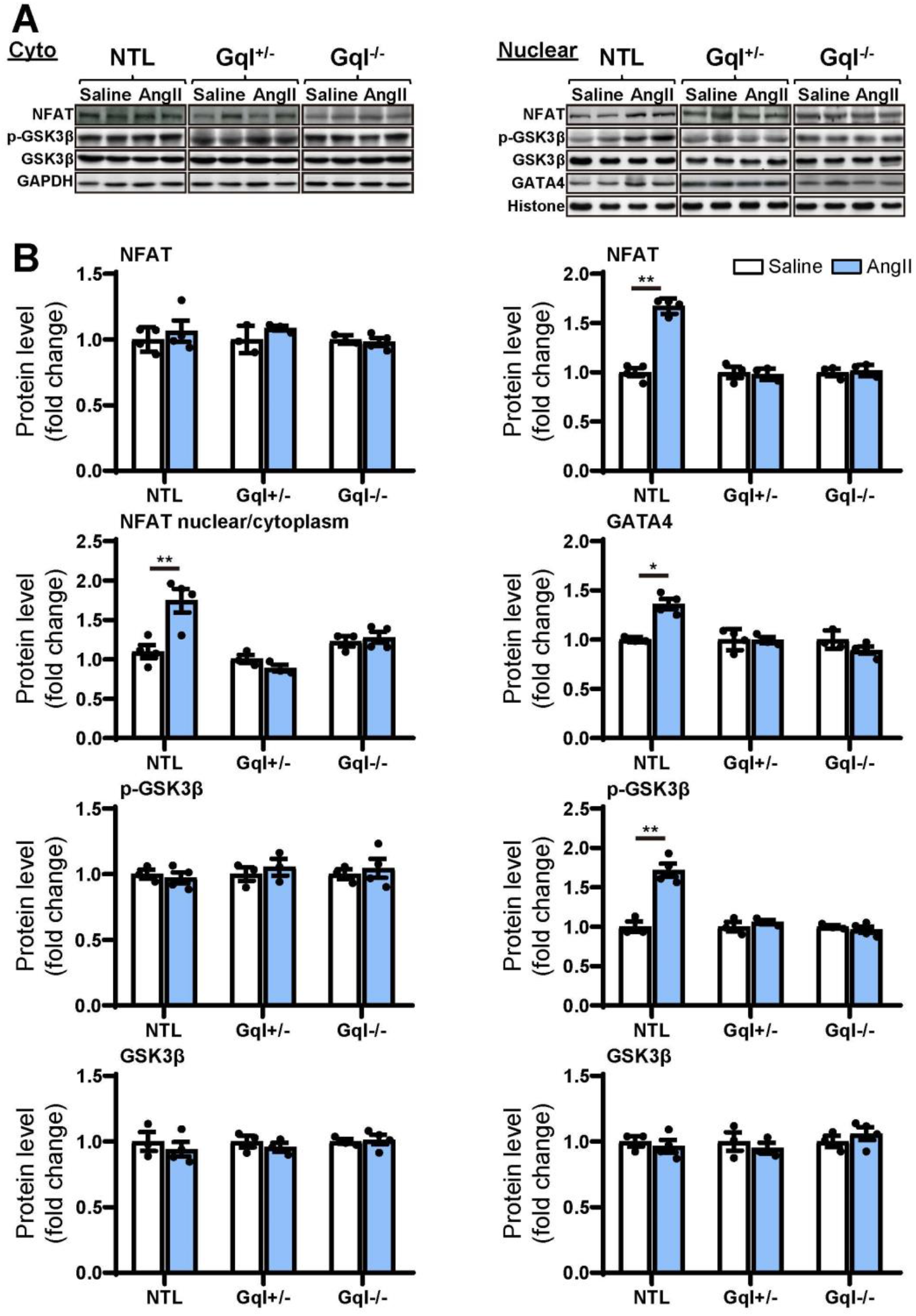
NFAT-GATA4 signalling in response to angiotensin II infusion after 48 hours. (A) Representative Western blots of NFATc4, phosphorylated glycogen synthase kinase-3 beta (GSK3β) and total GSK3β protein expression in the cytoplasm (upper left panel) and nucleus (upper right panel). GATA4 blots are shown for the nucleus. (B) Cytoplasmic and nuclear data are normalised with GAPDH and histone, respectively (bottom panels). All Western blot runs were performed in duplicate or triplicate. Results are presented as the mean value of 5 hearts per group ± SEM. \**p*<0.05, \*\**p*<0.01, vs. saline treated controls.

Importantly, all of these manifestations of the activation of the calcineurin-NFAT pathway with AngII infusion in NTL mice were completely absent in both the GqI^+/-^ and the GqI^-/-^ mice (Figure 4), consistent with the inhibition of the hypertrophic response to AngII in the transgenic hearts (Figure 1A).

### TAC does not activate the calcineurin-NFAT hypertrophic signaling pathway

In contrast to the effects of AngII infusion, TAC for 48 hours did not cause any significant change in the levels of NFATc4, GATA4, total GSK3β or phosphorylated GSK3β in the nucleus or the cytoplasm in NTL hearts, nor in GqI^+/-^ or GqI^-/-^ hearts (Figure 5), indicating that TAC does not activate the calcineurin-NFAT hypertrophic signaling pathway.

**Figure 5:**
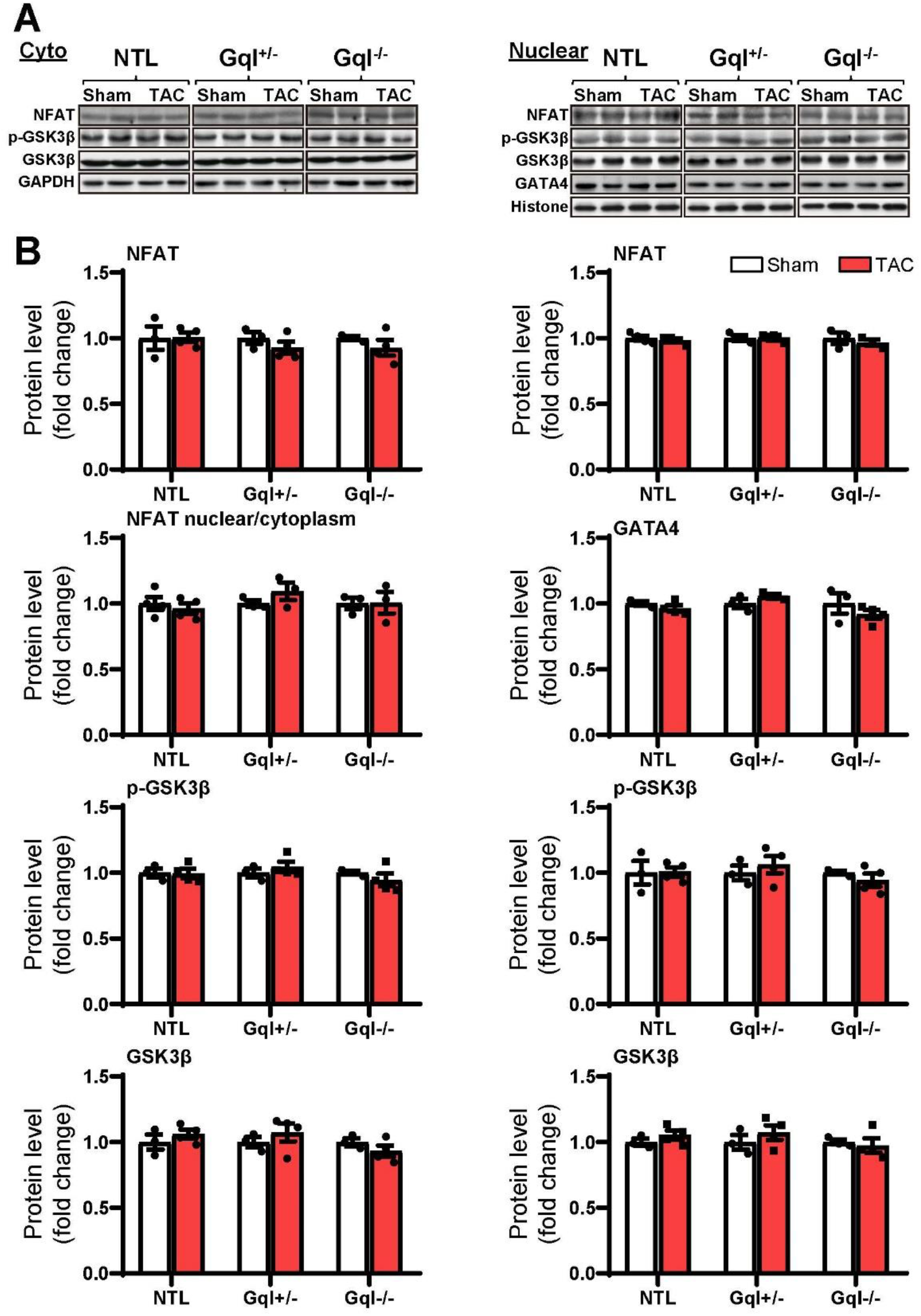
NFAT-GATA4 signalling in response to TAC after 48 hours. (A) Representative Western blots of NFATc4, phosphorylated glycogen synthase kinase-3 beta (GSK3β) and total GSK3β protein expression in the cytoplasm (upper left panel) and nucleus (upper right panel). GATA4 blots are shown for the nucleus. (B) Cytoplasmic and nuclear data are normalised with GAPDH and histone, respectively (bottom panels). All Western blot runs were performed in duplicate or triplicate. Results are presented as the mean value of 5 hearts per group ± SEM. None of the statistical comparisons between TAC mice and sham-operated controls were significant.

### Angiotensin II does not activate the CaMKII-HDAC4-MEF2 signaling pathway

The infusion of AngII for 48 hours did not cause any significant change in the levels of CaMKIIδ, total HDAC4 or phosphorylated HDAC4 (p-HDAC4) in the nucleus or the cytoplasm in NTL hearts, nor in GqI^+/-^ or GqI^-/-^ hearts (Figure 6). Nor was there any change in nuclear levels of MEF2A with AngII infusion in NTL or transgenic hearts. These findings indicate that AngII does not activate the CaMKII-HDAC4-MEF2 hypertrophic signaling pathway.

**Figure 6:**
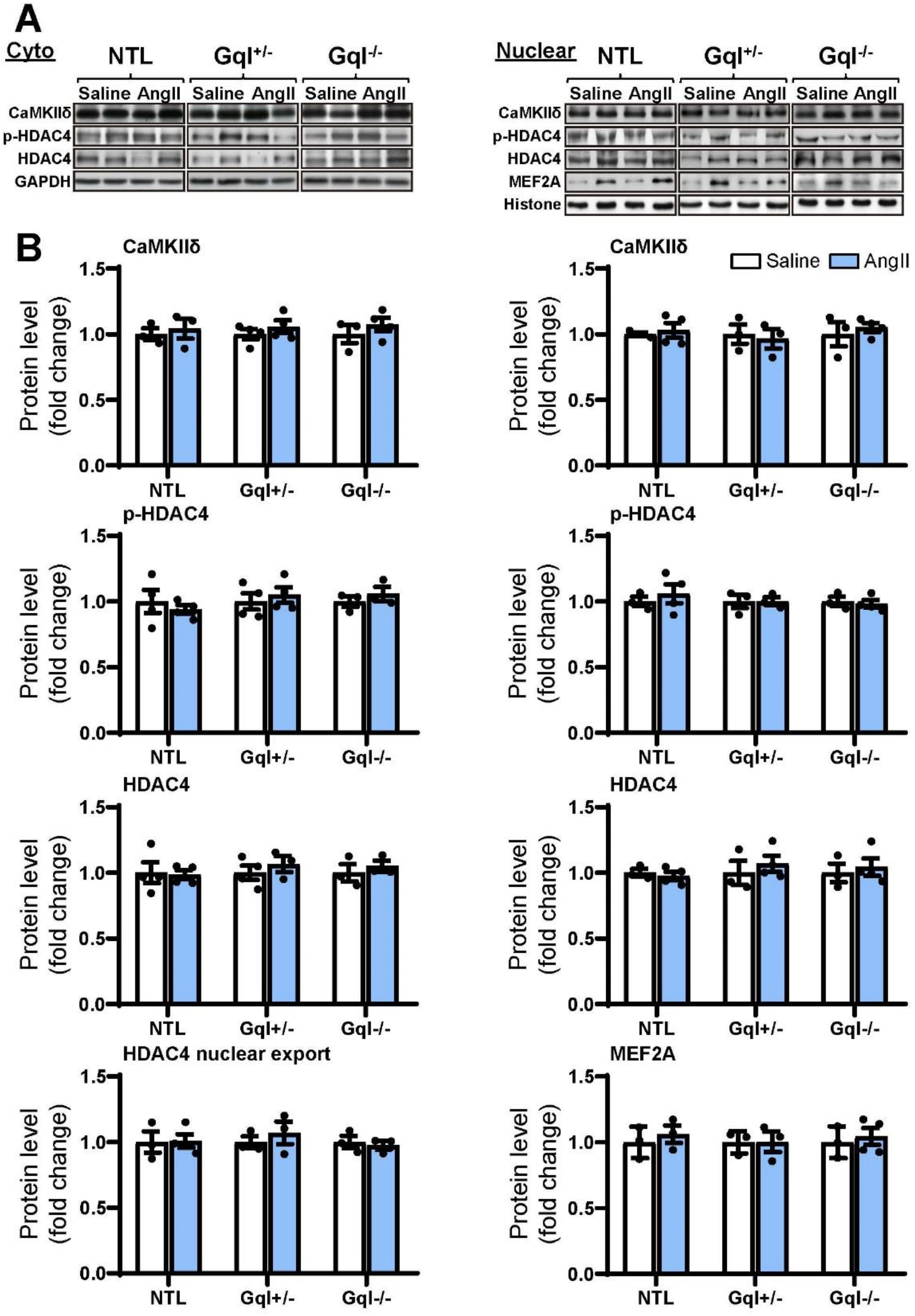
CaMKIIδ, HDAC4 and MEF2A signalling in response to angiotensin II after 48 hours. (A) Representative Western blots of CaMKIIδ, phosphorylated HDAC4 and total HDAC4 protein expression in the cytoplasm (upper left panel) and in the nucleus (upper right panel). MEF2A blots are shown for the nucleus. (B) Cytoplasmic and nuclear data are normalized with GAPDH and histone, respectively (bottom panels). The HDAC4 cytoplasmic/nuclear (cyto/nuclear) ratio was used as an indicator of HDAC4 nuclear export. All Western blot runs were performed in duplicate or triplicate. Results are presented as the mean value of 5 hearts per group ± SEM. None of the statistical comparisons between mice receiving AngII and saline were significant.

### TAC activates the CaMKII-HDAC4-MEF2 hypertrophic signaling pathway, and this activation is not inhibited by inhibition of cardiac Gq-coupled receptors

NTL hearts subjected to TAC for 48 hours exhibited a significant increase in the levels of CaMKIIδ in both the cytoplasm and the nucleus (both *p*<0.01, Figure 7). This TAC-induced increase in CaMKIIδ in NTL hearts was accompanied by increased levels of both total HDAC4 (p<0.05) and p-HDAC4 (p<0.01) in the cytoplasm, but there was no change in the nuclear HDAC4 or p-HDAC4 levels. Consequently, there was a significant increase in the cytoplasmic/nuclear ratio of HDAC4 (p<0.05) in NTL hearts, indicating increased nuclear export of HDAC4, and this was associated with a significant increase in the nuclear levels of MEF2A (p<0.001). These findings indicate that TAC activates the CaMKII-HDAC4-MEF2 hypertrophic pathway, resulting in nuclear export of HDAC4, relieving its repression of MEF2A.

**Figure 7:**
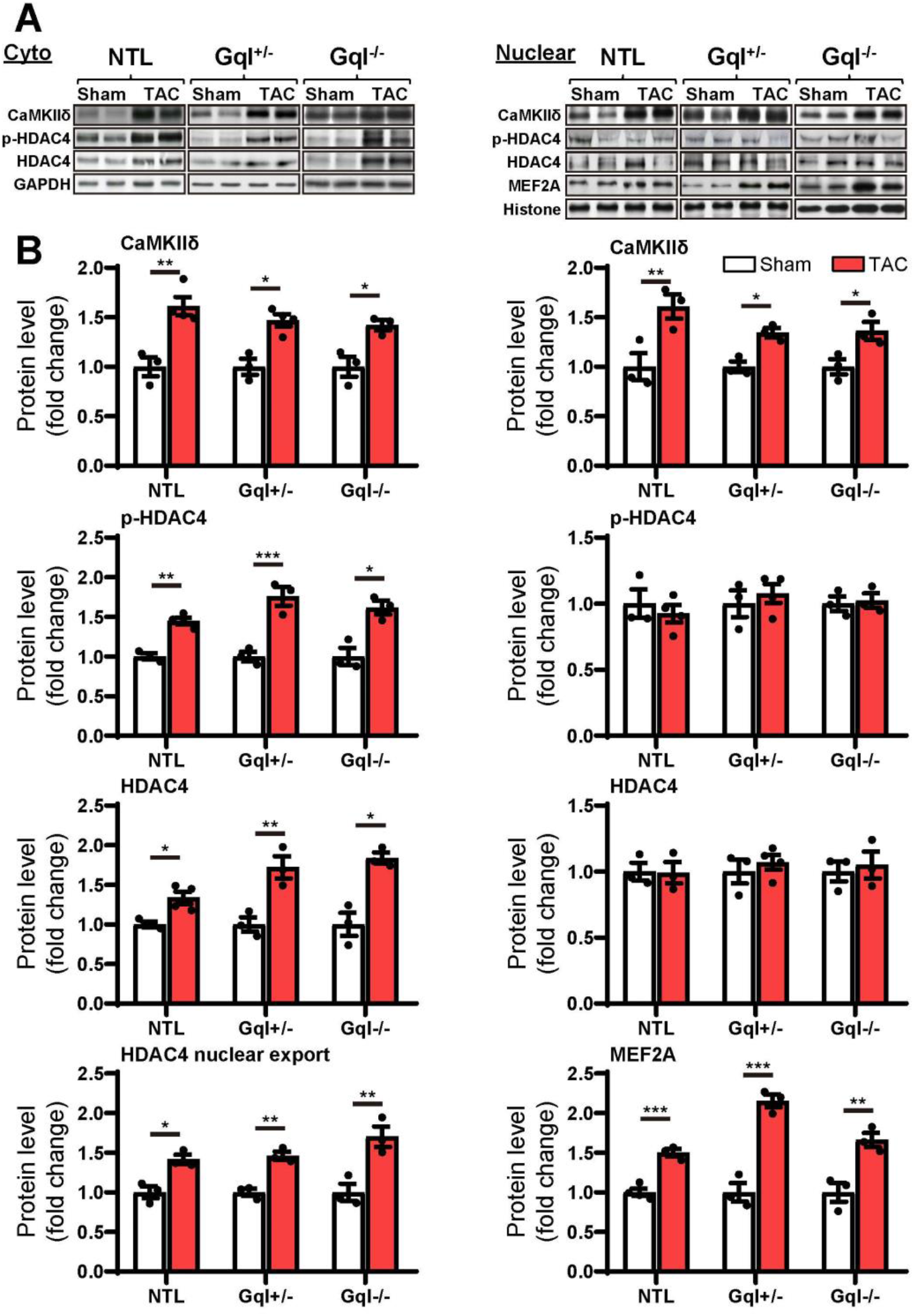
CaMKIIδ, HDAC4 and MEF2A signalling in response to TAC after 48 hours. (A) Representative Western blots of CaMKIIδ, phosphorylated (p-) HDAC4 and total HDAC4 protein expression in the cytoplasm (upper left panel) and in the nucleus (upper right panel). MEF2A blots are shown for the nucleus. (B) Cytoplasmic and nuclear data are normalized with GAPDH and histone, respectively (bottom panels). The HDAC4 cytoplasmic/nuclear (cyto/nuclear) ratio was used as an indicator of HDAC4 nuclear export. All Western blot runs were performed in duplicate or triplicate. Results are presented as the mean value of 5 hearts per group ± SEM. \**p*<0.05, \*\**p*<0.01, \*\*\**p*<0.001 vs. sham-operated controls.

Importantly, all of these manifestations of the activation of the CaMKII-HDAC4-MEF2 hypertrophic pathway with TAC in NTL mice were also present in the GqI^+/-^ and the GqI^-/-^ mice, indicating that inhibition of cardiac Gq-coupled receptors has no effect on the activation of the CaMKII-HDAC4-MEF2 pathway with TAC. These findings are consistent with the complete absence of any inhibition of the hypertrophic response to TAC in the transgenic GqI hearts (Figure 2A).

### MCIP1 is a marker of activation of both the calcineurin-NFAT and CaMKII-HDAC4-MEF2 hypertrophic signalling pathways

Because MCIP1 gene expression is activated downstream of the nuclear translocation of NFATc4, it is often measured as an *in vivo* marker of calcineurin-NFAT hypertrophic pathway activation. Gene expression of the MCIP1.4 isoform was significantly and similarly elevated 48 hours after *both* AngII infusion and TAC when compared with their respective saline infusion and sham controls (Table 1), despite the fact that calcineurin activation was evident with AngII infusion but not TAC (Figures 4 and 5). These findings indicate that MCIP1 gene expression is increased not only by calcineurin-NFAT pathway activation but also by activation of the CaMKII-HDAC4-MEF2 hypertrophic pathway.

**Table 1.**
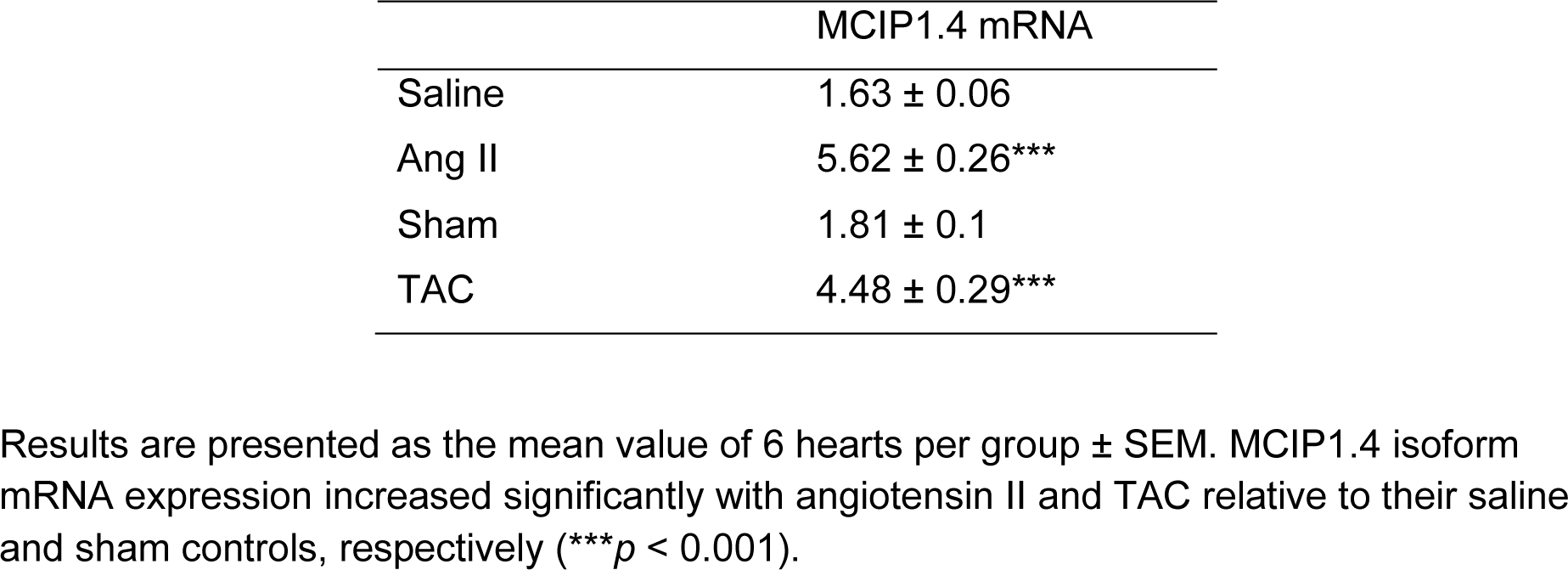
**MCIP1.4 isoform mRNA expression after angiotensin II (Ang II) or saline infusion for 48 hours and 48 hours after TAC or sham surgery in non-transgenic mice**.

|  | MCIP1.4 mRNA |
| --- | --- |
| Saline | 1.63 ± 0.06 |
| Ang II | 5.62 ± 0.26*** |
| Sham | 1.81 ± 0.1 |
| TAC | 4.48 ± 0.29*** |

Results are presented as the mean value of 6 hearts per group ± SEM. MCIP1.4 isoform mRNA expression increased significantly with angiotensin II and TAC relative to their saline and sham controls, respectively (\*\*\* $p < 0.001$ ).

## Discussion

### The role of cardiac Gq-coupled receptors in the induction of LVH secondary to pressure overload

The major finding of this investigation is that cardiac Gq-coupled receptors were not required for the induction of LVH after TAC, the most commonly employed experimental model of LVH secondary to LV pressure overload.

Although our results are very clear, and consistent with findings in mice with a global AT1a knockout after TAC (Harada et al., 1998b), they differ from those in the initial report of the phenotype of heterozygous GqI mice, which were reported to exhibit partial inhibition of TAC-induced LVH (Akhter et al., 1998). It is important to emphasise, therefore, that we took two steps in the current study to ensure that we achieved a robust inhibition of Gq. First, we bred both heterozygous (GqI^+/-^) and homozygous (GqI^-/-^) Gq-inhibited mice for this study. Second, the main purpose of the AngII infusion experiments in this study was to demonstrate that the Gq-inhibited mice used in this study did indeed exhibit effective Gq inhibition phenotypes. We demonstrated significant inhibition of AngII-induced LVH in mice with heterozygous Gq inhibition and complete inhibition of LVH in mice with homozygous Gq inhibition. Similarly, transgenic Gq inhibition prevented the increased gene expression of the hypertrophic markers ANP, BNP and α-SA with AngII infusion. Finally, the activation of the calcineurin-NFAT hypertrophic signalling pathway by AngII was inhibited by transgenic Gq inhibition. These results demonstrate clearly and in a ‘dose-dependent’ manner the effectiveness of the transgenic inhibition of Gq. The same results would be anticipated with other agonists of Gq-coupled receptors, including endothelin and catecholamines.

In contrast to the results obtained with AngII infusion, we showed that the transgenic inhibition of cardiac Gq-coupled receptors had *no effect* on the amount of TAC-induced LVH after 21 days, even with homozygous Gq inhibition. Consistent with this finding, transgenic Gq inhibition did not prevent the increased gene expression of ANP, BNP and α-SA with TAC. Finally, the activation of the CaMKII-HDAC4-MEF2 pathway by TAC was not inhibited by transgenic Gq inhibition.

Our results do not exclude a role for cardiac Gq-receptors in the induction of LVH in situations where a variety of humoral stimuli may be important. For example, the systemic renin-angiotensin system is activated in hypertension secondary to renal artery stenosis, and activation of Gq-coupled cardiac receptors by circulating AngII would be expected to play an important role in the induction of LVH in this situation. Similarly, elevated circulating catecholamine levels can induce LVH via their Gq-coupled α-adrenergic receptors, as can elevated endothelin levels acting via their Gq-coupled endothelin receptors.

### Differential role of two calcium-calmodulin dependent mechanisms in the induction of pressure overload LVH

The second major finding of this investigation is the remarkable *segregation* of the roles of two calcium-calmodulin dependent hypertrophic pathways in mediating LVH secondary to a humoral stimulus (AngII) acting via Gq-coupled receptors, on one hand, and LVH secondary to a mechanical increase in proximal aortic pressure (TAC), on the other. It is generally believed that because activation of both pathways is calcium-calmodulin dependent, they are activated together by a variety of pathological signals that increase intracellular calcium, and that there is significant cross-talk between these pathways (Zarain-Herzberg et al., 2011). These segregated hypertrophic pathways are shown schematically in Figure 8. Gq-coupled receptor activation by a ligand drives LVH via the calcium-calmodulin dependent activation of calcineurin, which dephosphorylates NFAT, resulting in increased NFAT translocation to the nucleus, inhibition of GSK3β-mediated export of NFAT from the nucleus, increased GATA4 and combinatorial interaction with MEF2 that initiates transcription of hypertrophic genes. On the other hand, a mechanical stimulus (TAC) drives LVH via calcium-calmodulin dependent activation of CaMKII, resulting in increased nuclear export of HDAC4, which relieves the repression of MEF2, permitting its combinatorial interaction with NFAT and GATA4 to initiate hypertrophic gene transcription.

**Figure 8:**
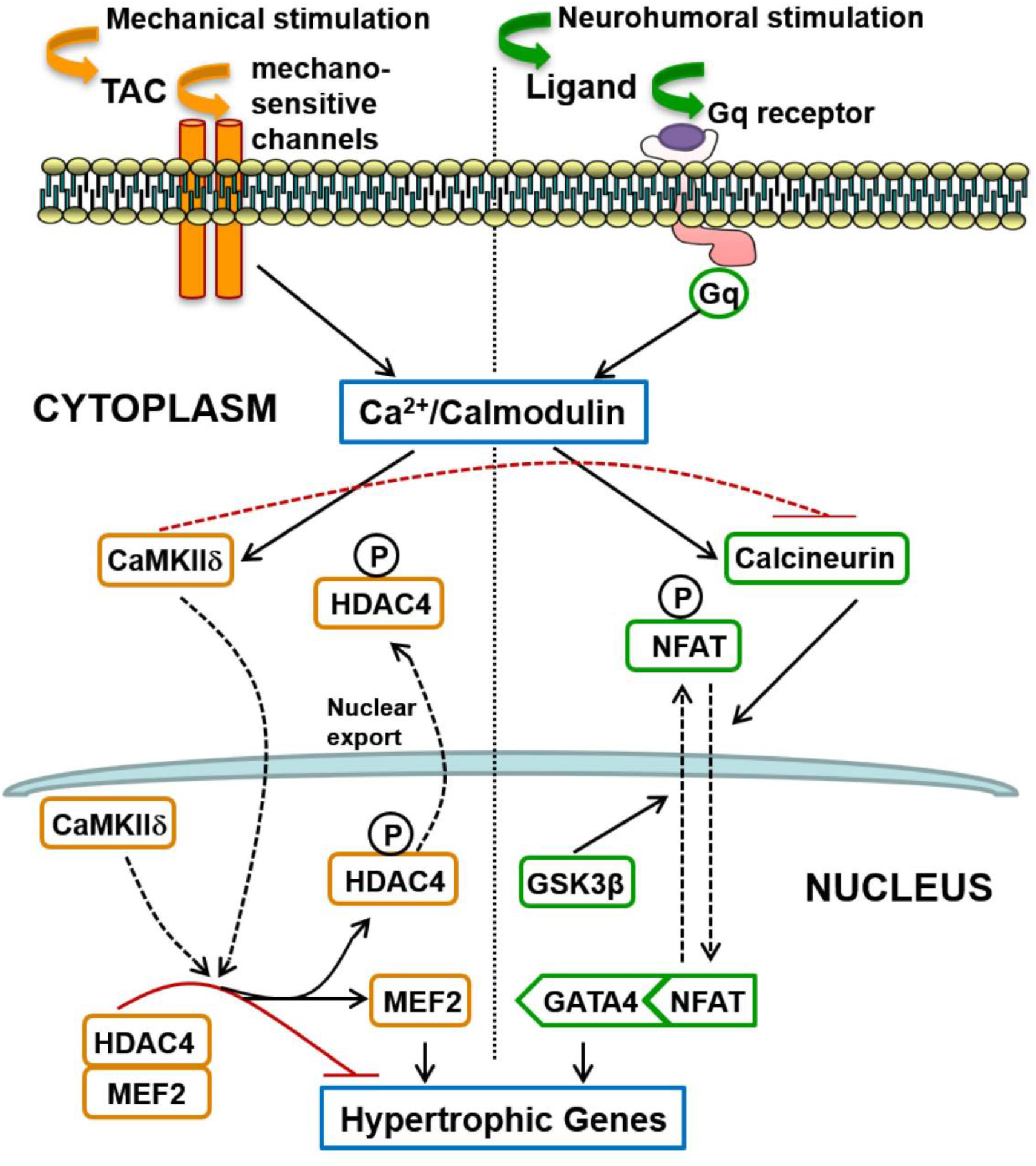
Schematic summary of two distinct signalling pathways involved in the induction of cardiac hypertrophy in response to pressure overload: (1) The calcineurin-NFAT pathway is activated by Gq-coupled receptors in response to neurohumoral stimulation, not only by AngII but possibly also by endothelin and by catecholamines (acting on alpha-adrenergic receptors); (2) Mechanical loading, such as transverse aortic constriction (TAC), does not activate the systemic renin-angiotensin system but induces hypertrophy mediated by CaMKII-HDAC4-MEF2 signalling that does not require Gq-coupled receptor activation, but is presumably activated by calcium entry controlled by mechano-sensitive channels. Differences in the mode of intracellular calcium elevation in response to different stimuli (mechanical versus neuro-humoral) may account for the differential activation of calcineurin versus CaMKII, but CaMKII also directly inhibits activation of calcineurin, as indicated by the broken red inhibitory path (see Discussion).

### MCIP1 is a marker of both calcineurin and CaMKII activation

The third important finding of this investigation is that MCIP1 gene expression is increased not only with calcineurin pathway activation, as expected, but also with CaMKII pathway activation (Table 1). This is a consequence of the convergence of these two distinct signalling pathways onto a common intra-nuclear complex to initiate hypertrophic transcription (Figure 8) and the fact that nuclear export of HDAC4 relieves the repression not only of MEF2 but also of NFAT (McKinsey, 2007;Backs et al., 2011). Derepression of both MEF2 and NFAT by CaMKII activation results in gene transcription downstream of intra-nuclear NFAT, including expression of MCIP1 (Figure 8). It follows that increased MCIP1 gene expression cannot be regarded as a specific marker of calcineurin activation. For the same reason, a similar caveat should apply to conclusions regarding calcineurin activation drawn from experiments based on the NFAT luciferase reporter (Wilkins et al., 2004;Molkentin, 2013).

### Mechanisms of differential activation of calcineurin and CaMKII

The interesting question is the mechanism of differential activation of calcineurin and CaMKII in response to TAC given that both depend on calcium-calmodulin activation. It is known that calcineurin activation requires a sustained increase in the resting intracellular calcium level (Dolmetsch et al., 1997). Sustained Gq-coupled receptor activation by its humoral ligand (angiotensin II, endothelin or catecholamines), with activation and increased expression of TRPC6 receptors (Eder and Molkentin, 2011), for example, could provide the mechanism for this tonic calcium signal. In contrast, CaMKII activation is more sensitive to high-frequency, high amplitude calcium oscillations (De Koninck and Schulman, 1998), and it is known that aortic constriction provides this type of calcium signal (Colella et al., 2008), possibly initiated directly by mechano-sensitive channels given our new findings regarding Gq (Figure 8), although the inititial mechano-transduction could then trigger secondary sources of calcium entry (see Introduction).

In addition, activated CaMKII has been shown to directly inhibit the regulatory subunit of calcineurin (Kreusser et al., 2014), providing an additional explanation for the differential signalling pathway activation with TAC. This direct inhibition of calcineurin by CaMKII is also consistent with our finding that Gq inhibition has no impact on the hypertrophic response to TAC despite the fact that TAC increases DAG, which indicates Gq activation (Niizeki et al., 2008).

While there is consensus that activation of the CaMKII-HDAC4-MEF2 pathway causes the adverse left ventricular remodelling and heart failure that is associated with TAC-induced LVH (Ling et al., 2009;Kreusser et al., 2014), it has been less clear to some that this pathway induces the hypertrophy *per se*. A CaMKIIδ knockout (KO) in the mouse heart was clearly shown to inhibit TAC-induced LVH in 2009 (Backs et al., 2009), but a subsequent study by others of a CaMKIIδ KO mouse in the same year showed no inhibition of LVH 2 weeks after TAC, despite protection from adverse remodelling and heart failure 6 weeks after TAC (Ling et al., 2009). In the latter study, however, there was clear inhibition of LVH with CaMKIIδ KO 6 weeks after TAC but this was interpreted as secondary to the protection from adverse LV remodelling and dilatation. A subsequent study of a double CaMKII isoform KO model presented a more complex picture of the hypertrophic response (Kreusser et al., 2014). In that latter study, a single KO of *either* CaMKIIδ or CaMKIIγ in the mouse heart did significantly inhibit the hypertrophic response to TAC (Kreusser et al., 2014), while a double KO of *both* CaMKII isoforms inhibited the adverse left ventricular remodelling and cardiac dysfunction but the hypertrophic response to TAC *increased* when compared with the single KO experiments, and was similar to control animals. This was explained by the loss of the direct inhibitory effect of CaMKII on calcineurin with the CaMKII double isoform KO (see above), so that the resultant increase in calcineurin activation explained the increased hypertrophy in the double KO animals when compared with the single KO animals (Kreusser et al., 2014).

It would be a mistake to infer from the CaMKII isoform double KO experiments above (Kreusser et al., 2014), however, that calcineurin activation induces the hypertrophic response to TAC in the normal heart while CaMKII activation induces only the adverse remodelling and heart failure. If that were the case, the CaMKII single isoform KO experiments would not have demonstrated the very significant reduction of LVH with TAC when compared with control animals (Backs et al., 2009;Kreusser et al., 2014). The alternative CaMKII isoform was shown to compensate for the loss of the direct inhibitory effect of the missing CaMKII isoform on calcineurin in the single isoform KO animals (Kreusser et al., 2014), with the consequence that the hypertrophic response to TAC in single KO animals should have equalled that in control animals unless CaMKII activation is pro-hypertrophic *per se*. Our results in the present study are consistent with this interpretation of the pro-hypertrophic effect of CaMKII activation. In the normal adult mouse heart, we observed activation of the CaMKII-HDAC4-MEF2 pathway preceding TAC-induced LVH, but there was no evidence of calcineurin activation.

It is relevant to note here also the recent observation that gene therapy with the N-terminal proteolytic fragment of HDAC4, HDAC4-NT, overcomes CaMKII-induced cytoplasmic accumulation of HDAC4 and subsequent MEF2 activation after TAC, inhibiting not just the adverse LV remodelling after TAC but also the LVH (Lehmann et al., 2018), which clearly indicates a pro-hypertrophic role of CaMKII activation via HDAC4, independent of calcineurin. In addition, the significant reduction in HDAC4-NT levels observed after TAC may play a significant role in TAC-induced LVH and adverse LV remodelling (Lehmann et al., 2018).

### Previous studies of calcineurin activation as the cause of pressure overload LVH

The large body of conflicting data concerning the importance of calcineurin activation in pressure overload LVH has been reviewed extensively elsewhere (Molkentin, 2000;2013). In addition to the previously acknowledged limitations of early studies based on pharmacological inhibitors of calcineurin, such as cyclosporin A and FK506, and the limitations of early calcineurin activity assays (Molkentin, 2000), it is important to consider the experimental pressure overload model employed in evaluating the results of previous studies. Most studies based on pressure overload induced by abdominal aortic constriction (AAC), for example, have reported that calcineurin-NFAT pathway activation accounts for the LVH observed, and this is consistent with our current results obtained with AngII infusion because AAC activates the systemic renin-angiotensin system (RAS) when the aorta is constricted between the renal arteries (Wiesner et al., 1997). For example, consistent with our AngII data in GqI mice, mice with condition inactivation of Gαq/Gα11 in cardiac myocytes were shown not to develop LVH secondary to pressure overload induced by AAC (Wettschureck et al., 2001). When supra-renal AAC is used, however, there is little or no evidence of RAS activation (Nicks et al., 2020), which is consistent with the absence of any reduction in the amount of LVH induced by this model in mice with a global AT1a receptor knockout (Harada et al., 1998a). Of those studies based on TAC, which does not cause systemic RAS activation and has become the dominant experimental model of pressure overload, the results with regard to calcineurin activation are much more variable (Frey and Olson, 2003;Molkentin, 2013). Many of these TAC studies have relied on MCIP1 or the NFAT luciferase reporter as the only in vivo evidence for calcineurin activation with TAC (Wilkins et al., 2004;Molkentin, 2013). For the reasons noted above, neither of these indices is a specific marker of calcineurin activation because CaMKII activation causes derepression of intra-nuclear NFAT as well as MEF2. The only incontrovertible evidence of calcineurin-NFAT pathway activation is direct measurement of nuclear translocation of NFAT based on measurements of the nuclear and cytoplasmic NFAT fractions (Molkentin, 2013), as was done in our current study. We observed similar elevations of MCIP1 with calcineurin activation alone and with CaMKII activation in the absence of calcineurin activation (Table 1).

Transgenic inhibition/inactivation/overexpression of calcineurin by a variety of approaches was important in highlighting the potential role of calcineurin in cardiac growth and hypertrophy, but there are some limitations to these methods (Molkentin, 2013). For example, mice with an inducible loss of all cardiac myocyte calcineurin activity showed early lethality, reduced myocyte proliferation rates with fewer myocytes, greater myocyte apoptosis rates with acute pressure overload, and greater cell death and injury after ischaemia-reperfusion injury (Molkentin, 2013). Similarly, mice with transgenic overexpression of the calcineurin inhibitory domains of Cain or A-kinase anchoring protein 79 exhibited reduced heart weights with thinner left ventricular walls and reduced myofibrillar cross-sectional areas relative to controls (Molkentin, 2013). These findings indicate that calcineurin is important for normal myocyte development and functioning, so that transgenic models that inhibit calcineurin early in life may exhibit a non-specific impairment of growth and hypertrophic responses to a variety of stimuli. Such models may be unreliable indicators of the specific role of calcineurin activation in response to pressure overload in the normal adult heart. Similarly, much of the evidence that calcineurin induces pathological cardiac hypertrophy with adverse remodelling and heart failure comes from studies that used forced overexpression of a truncated CnA construct that was deficient in the regulatory domain that CaMKII phosphorylates to inhibit calcineurin (Molkentin, 2013). Due to their lack of calcineurin inhibition with CaMKII activation, such models likely overstate the calcineurin-related hypertrophic response and the adverse remodelling response to various stimuli, including TAC (Kreusser et al., 2014). One advantage of the data presented here regarding the role of calcineurin in pressure overload hypertrophy is that there was no genetic manipulation of the calcineurin pathway.

### Clinical Implications

Our results may help explain why angiotensin converting enzyme inhibitors and angiotensin receptor blockers, although effective and frequently prescribed anti-hypertensive therapies, have not been shown to be more effective in reducing the morbidity and mortality of hypertensive heart disease than less specific anti-hypertensive agents, including diuretics (Psaty et al., 2003;Xue et al., 2015). This evidence suggests that for an equivalent blood pressure lowering effect, there is no convincing difference between anti-hypertensive therapies despite the putative anti-hypertrophic effects of newer and more expensive agents that target angiotensin and its receptors. Our results suggest that if the therapeutic aim is to prevent pressure-overload induce LVH and adverse remodelling, CaMKII or its upstream activators may be more effective targets for the development of new therapies.

## Abbreviations

LVH: left ventricular hypertrophy
AngII: angiotensin II
NFAT: nuclear factor of activated T-cells
CaMKII: calmodulin kinase II
HDAC4: histone deacetylase 4
MEF2: myocyte enhancer factor 2
TAC: transverse aortic arch constriction
LVW/BW: left ventricular weight normalized to body weight
LVW/TL: left ventricular weight normalized to tibia length
ANP: atrial natriuretic peptide
BNP: B-type natriuretic peptide
α-SA: α-skeletal actin

## ETHICS STATEMENT

All experimental procedures were approved by the Garvan/St Vincent’s Hospital Animal Ethics Committee, in accordance with the guidelines of the Australian Code of Practice for the Care and Use of Animals for Scientific Purposes.

## AUTHOR CONTRIBUTIONS

ZY and HG, joint first author. ZY, HG, JW, YD and SK conducted experiments and analysed results. DF, BM and RG contributed to experimental protocols and critically reviewed the experimental results and the manuscript. MF conceived and designed the experiments, and supervised their conduct. ZY and MF wrote the manuscript.

